# Sleep reactivation shapes memories of event and its temporal structure

**DOI:** 10.64898/2026.10.20.760365

**Authors:** Yuqi Zhang, Danni Chen, Tao Xia, Tianqi Di, Ziqing Yao, Winny W. Y. Yue, Zexuan Mu, Xiaoqing Hu

## Abstract

Episodic memory provides a temporal scaffold that organizes past experiences according to their sequential occurrence. While memories are reactivated and consolidated during sleep, it remains unknown how exogenous reactivation reshapes memories of complex events—particularly the temporal structure that binds individual elements across time. In this preregistered study, 35 participants encoded neutral and aversive narratives composed of image sequences, with auditory context shifts defining event boundaries. During post-encoding slow-wave sleep, closed-loop targeted memory reactivation (TMR) delivered boundary-associated auditory cues time-locked to the up-state of slow oscillations. Reactivation led to faster, more confident, and more liberal recognition responses for cued than uncued memories. However, reactivation exerted a time-dependent effect on temporal-order memory, impairing it immediately after sleep but attenuating forgetting one week later. This behavioral dissociation was paralleled at the neural level: stronger cue-elicited theta activity predicted superior recognition of neutral—but not aversive—event content, whereas greater theta responses were associated with poorer temporal-order memory for the same events. Together, these findings suggest that sleep reactivation does not uniformly strengthen episodic memories. Rather, it selectively prioritizes access to event content while reorganizing temporal relations among event elements, revealing distinct, time-dependent consequences of sleep reactivation for different components of complex experiences.

## Introduction

Episodic memory enables us to mentally reconstruct and relive past experiences, constituting a cornerstone of human cognition (Tulving, 2002). Rather than being replicas of the past, episodic memories are constructive and continuously transformed in both content and temporal structure (Dudai, 2012; Schacter, 2012). An exotic retreat may later be remembered as a series of discrete highlights—a stunning sunset or an unexpected injury—while memories of intervening episodes and moments fade and become blurred. Notably, memory transformations are not passively driven by the passage of time but are actively shaped during post-encoding sleep, when daytime experiences are reactivated and consolidated (Lutz et al., 2026; Tarder-Stoll et al., 2026). Yet how sleep reactivation may differentially reshape the content and temporal structure of complex events, and thus impact long-term stabilization and reconstruction, remains poorly understood.

One fundamental principle underlying the formation and reconstruction of episodic memory is event segmentation, whereby ongoing experience is parsed into discrete subevents at moments of contextual change, i.e., event boundaries (Bailey & Smith, 2024; Clewett et al., 2019; Zacks et al., 2007). Boundaries reorganize what is retained and how memories are temporally organized. Specifically, boundaries enhance memories of items adjacent to them but impair those further away (Heusser et al., 2018; Swallow et al., 2009). Meanwhile, boundaries retained memories of temporal order within a segment but impaired those across boundaries (Clewett et al., 2019; DuBrow & Davachi, 2016). Neuroimaging evidence further shows that event boundaries trigger abrupt neural state transitions, accompanied by reinstatement of the pre-boundary event representations (Davachi & DuBrow, 2015; Hahamy et al., 2023; Silva et al., 2025). Such boundary-related neural dynamics are thought to stabilize boundary-proximal event content while weakening temporal memories of sub-events across boundaries (Clewett et al., 2024; Sols et al., 2017).

Event boundaries, however, are not restricted to contextual changes; internal affective shifts can similarly segment ongoing experience (Clewett et al., 2019). Mounting evidence has established that emotional salience enhances memory for affect-laden event content (LaBar & Cabeza, 2006; Payne & Kensinger, 2010). Beyond event content, emotion also shapes memories of temporal structure: positive emotional shifts strengthen temporal-order memory between adjacent subevents, whereas negative shifts promote temporal separation between subevents (McClay et al., 2023; Petrucci & Palombo, 2021; J. Wang et al., 2022). Notably, emotional salience can retroactively enhance neutral memories that are temporally adjacent or conceptually related to an emotionally salient event (Clewett & McClay, 2025; Dunsmoor et al., 2015, 2018; Rouhani et al., 2023). Together, these findings establish emotional shifts as boundaries that organize event memories, However, whether emotional boundaries continue to bias the offline consolidation of event content and its temporal structure remains unknown.

During sleep, memories of prior experiences are reactivated and gradually transformed through cross-region, systems-level consolidation (Brodt et al., 2023; Klinzing et al., 2019). In rodents, hippocampal neural ensembles associated with daytime experience (e.g., navigating a maze) are sequentially reactivated during sleep, i.e., memory replay (Huelin Gorriz et al., 2023; Ji & Wilson, 2007; K. Liu et al., 2024). By recapitulating prior experiences in a temporally compressed form, replay supports memory consolidation (Buzsáki, 2015; Z. S. Chen & Wilson, 2023; Foster, 2017). More critically, the inherently sequential nature of replay provides a plausible mechanism for preserving the temporal structure of continuous experiences (J. Zhang et al., 2025). Converging evidence in humans similarly links offline reactivation of individual memory elements to subsequent retention (J. Liu et al., 2023; Schapiro et al., 2018; Schreiner et al., 2021), whereas sequential replay during awake rest reflects the ordered structure of learned experiences (Y. Liu et al., 2019; Schuck & Niv, 2019). However, these findings largely stem from paired associates or short sequences, leaving open whether these principles during sleep extend to temporally extended, structured events (Antony & Schechtman, 2023).

For emotional events, sleep-dependent consolidation may differentially affect memories of event content and temporal structure. Emotional salience can shape which memories are prioritized for offline reactivation, potentially through coordinated amygdala–hippocampal processing (Cabrera et al., 2024; Girardeau et al., 2017; Sterpenich et al., 2021). Regarding temporal memories, sleep preserves the sequential order of autobiographical memories (e.g., an art tour) while perceptual details faded over time (Diamond et al., 2025; Drosopoulos et al., 2007; Lerner & Gluck, 2019). Given that emotional arousal can alter how successive moments are temporally integrated or separated during encoding (McClay et al., 2023; Petrucci & Palombo, 2021), an unanswered question is whether subsequent sleep reactivation stabilizes or reorganizes the temporal structure of an emotional experience. Addressing this question requires moving beyond isolated items to consider how reactivation operates within emotionally structured events. More intriguingly, recent evidence suggests that the benefits of sleep reactivation can generalize beyond directly reactivated memories to their contextual associates (Schechtman et al., 2023a, 2023b), raising the possibility that reactivation propagates through the conceptual and temporal structure of an event rather than benefiting cued items in isolation. Yet whether such effects extend to naturalistic events organized by emotional boundaries and temporal sequences remains untested. Thus, the current study bridges this gap by delineating how reactivation spreads within and across emotional boundaries to shape memories of event content and its temporal structure.

To causally investigate how sleep reactivation shapes memories of emotional events, we re-presented memory-associated acoustic cues to sleeping participants to experimentally induce memory reactivation, i.e., targeted memory reactivation (TMR) (Carbone & Diekelmann, 2024; Hu et al., 2020). Specifically, participants encoded image sequences in which a shift in the auditory context signaled an event boundary, dividing each event into a neutral pre-boundary subevent followed by either an aversive or a neutral post-boundary subevent. During subsequent slow-wave sleep, we employed closed-loop stimulation to deliver boundary cues timed-locked to the up-state of ongoing slow oscillations to reactivate the associated memories. Because sleep reactivation may modulate the affective impact of emotional memories, albeit with mixed results (Hauner et al., 2013; Hu et al., 2020; Hutchison et al., 2021; Recher et al., 2024), we also examined participants’ emotional responses to event memories. This experiment thus allowed us to examine whether and how reactivating neutral and aversive event memories during sleep would modulate content memory (what happened), temporal memory (when it happened), and memory-specific affective responses (Fig. 1).

**Fig. 1.**
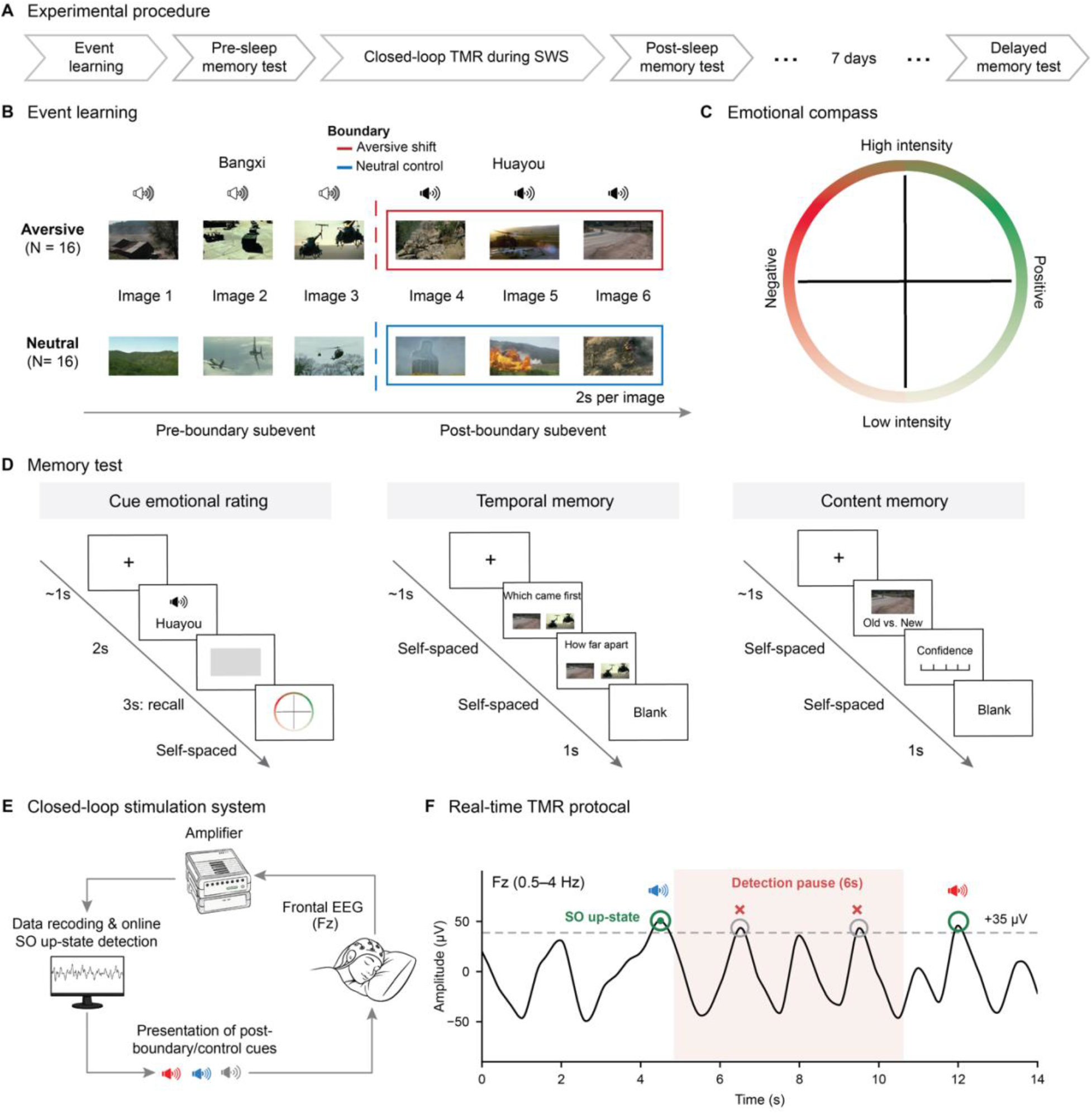
Experimental design. **(A) Procedure**. Participants visited the laboratory twice, separated by one week. The first visit comprised encoding, pre- and post-sleep memory tests, and closed-loop TMR during post-encoding slow-wave sleep. One week later, participants returned for a delayed memory test. **(B) Event learning.** Participants learned 32 events, each consisting of six images extracted from a short movie clip (all clip images depict scenes without identifiable individuals and are shown for illustrative purposes only). Each event was paired with two pseudowords that divided the six images into two three-image subevents, with the onset of the second pseudoword marking the event boundary. Participants were instructed to mentally construct a coherent story and memorize both the content and the temporal order of each event. **(C) Emotional compass.** Participants rated the valence and arousal of each cue on a two-dimensional compass. A planchette (green/red circle) tracked the cursor position and changed color according to its placement. **(D) Memory tests.** Before sleep, after sleep, and at the one-week delay, participants completed tests assessing emotional responses to the cues and temporal memory. Content memory was assessed via a recognition task administered only at the delayed session, given ceiling-level encoding performance. **(E-F) Closed-loop TMR protocol.** The second pseudoword of each event (i.e., the post-boundary cue) served as an auditory memory cue and was re-presented during slow-wave sleep, time-locked to the up-state of ongoing slow oscillations recorded at Fz. Half of the events were cued; novel sounds unassociated with any learned event served as control cues.

We preregistered the hypotheses that reactivation would: (1) enhance content memory for cued post-boundary subevents; (2) strengthen temporal memory for neutral sequences but disrupt temporal memory for aversive sequences; and (3) attenuate emotional responses to cued aversive subevents. In addition, because cued post-boundary subevents were embedded within broader event sequences, we further examined whether reactivating post-boundary memories would generalize to impact pre-boundary memories and the pre- and post-boundary temporal memories (see https://osf.io/5afw8/overview?view_only=0db0a541881f4ca984b0b40b69682bdf and Supplementary Table S1 for preregistration; deviations from the preregistered analyses are summarized in Table S2).

## Results

### Preregistered analyses on content memory

We used signal detection measures to quantify sensivity and response biases derived from old/new recognition task administered one week post-sleep. A 2 (TMR: cued vs. uncued) × 2 (event valence: aversive vs. neutral) × 2 (boundary position: pre vs. post-boundary) repeated-measures ANOVA on recognition sensitivity (*d′*) revealed no significant TMR effect, *F*(1, 30) = 1.33, *p* = .258, 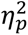 = .04, despite numerically higher *d′* in the cued condition. This null effect likely reflects near ceiling recognition even after one-week delay (d′ ≥ 3.46, hit rate ≥ 0.91), potentially limiting sensitivity to detect cueing-related benefits. However, cueing shifted response criteria toward more liberal (i.e., a greater tendency to respond "old") than the uncued condition, *F*(1, 30) = 4.41, *p* = .044, 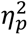 = .17. No other effects were significant (all *ps* ≥ .142). Separate repeated-measures ANOVAs on log-transformed reaction time and confidence rating for old items further indicated that cueing produced faster and more confident responses, *F*(1, 30) = 6.02, *p* = .020, 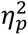 = .17 and *F*(1, 30) = 4.67, *p* = .039, 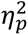 = .13 (Fig. 2A). Valence nor boundary position did not interact with TMR in modulating these effects (all *ps* > .40). Collectively, these findings suggest that TMR facilitated access to content memory rather than recognition sensitivity, yielding faster, more confident, and more liberal recognition responses.

**Fig. 2.**
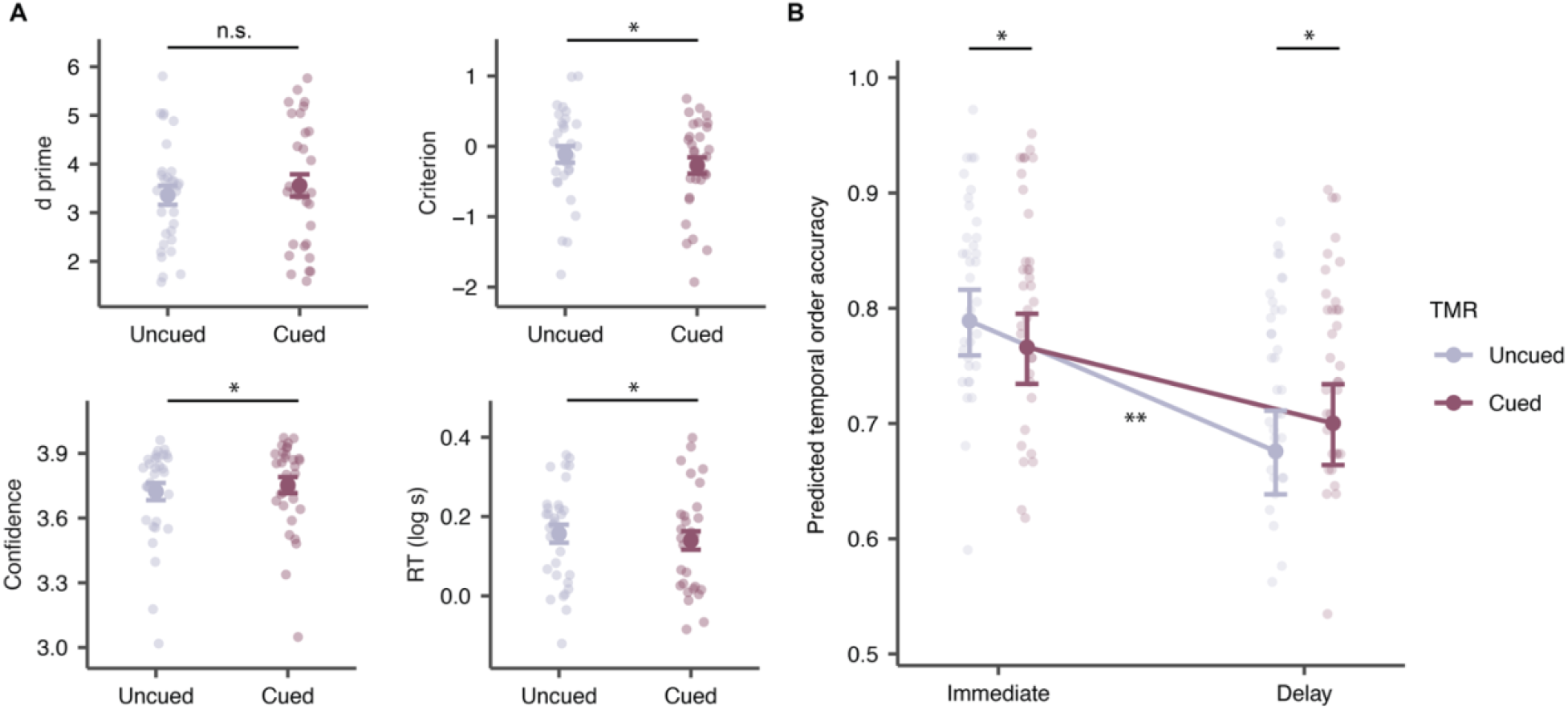
(A) No significant TMR effect was observed on recognition sensitivity, although a numerical increase was evident. By contrast, TMR significantly reduced response criterion, increased confidence ratings, and accelerated responses during content-memory judgments. (B) TMR impaired temporal memory immediately after sleep but attenuated its decline across the delay. Dots represent individual participants; \**p* < .05; n.s., not significant.

### Preregistered analyses on event temporal memory

Temporal-order accuracy and temporal distance estimates were analyzed using trial-level generalized linear mixed-effects models (GLMMs) and LMMs, respectively, with TMR (cued vs. uncued), event valence (aversive vs. neutral), boundary position (pre-, post-, and cross-boundary), test time (immediate vs. delayed), and all their interactions as fixed effects. Pre-sleep baseline ratings were included as a covariate, with random intercepts for participant and event. For temporal-order memory accuracy, although TMR effect was not significant, *χ*²(1) = 0.05, *p* = .816, a significant TMR × test time interaction emerged, *χ*²(1) = 9.91, *p* = .002. Follow-up simple effects analyses revealed that the direction of the TMR effect reversed from the immediate to the delayed tests: reactivation reduced temporal-order accuracy at the immediate test (76.6% vs. 78.9%, *OR* = 0.88, *z* = -2.26, *p* = .024); whereas at the delayed test, cued events showed higher accuracy than uncued events (70.0% vs. 67.6%, *OR* = 1.12, *z* = 2.17, *p* = .030, Fig. 2B), suggesting that cueing reduced forgetting across the retention interval. In addition to the interaction, the model revealed significant main effects of event valence, *χ*²(1) = 39.60, *p* < .001, boundary position, *χ*²(2) = 203.16, *p* < .001, and test time, *χ*²(1) = 139.38, *p* < .001. Specifically, accuracies were significantly higher for aversive than for neutral events (78.5% vs. 67.9%), and were substantially higher for cross-boundary pairs than for pre- or post-boundary pairs (81.0% vs. 69.5% and 68.8%, respectively), and declined from the immediate to the delayed test (77.8% vs. 68.8%). In contrast, no significant TMR-related effects were observed for temporal distance estimates (all *ps* > .146).

### Preregistered analyses on event-related emotional responses

Trial-level LMMs showed no significant TMR effect nor TMR-related interactions on valence and arousal ratings (all *ps* > .340, see Supplementary Data S1 for details).

### Preregistered EEG and EEG-behavior analyses

#### Closed-loop cue-elicited EEG responses during slow-wave sleep

We first verified the accuracies of the closed-loop SO detection and cueing procedure by analyzing cue-elicited event-related brain potentials (ERPs) and the phase distribution of cue delivery during slow-wave sleep (Fig. 3A-B). Offline validation verified that on average, 93.3% of cues (SD = 2.6%, range = 86.9– 99.4%) were delivered during the SO up-state, demonstrating successful closed-loop SO detection and stimulation. We next analyzed time-frequency-resolved EEG responses elicited by memory cues (aversive and neutral) and memory-unrelated control cues. EEG responses were averaged across frontocentral electrodes (F1/2, FC1/2, C1/C2, Fz, FCz and Cz) and compared between conditions using cluster-based permutation tests across time-frequency points.

**Fig. 3.**
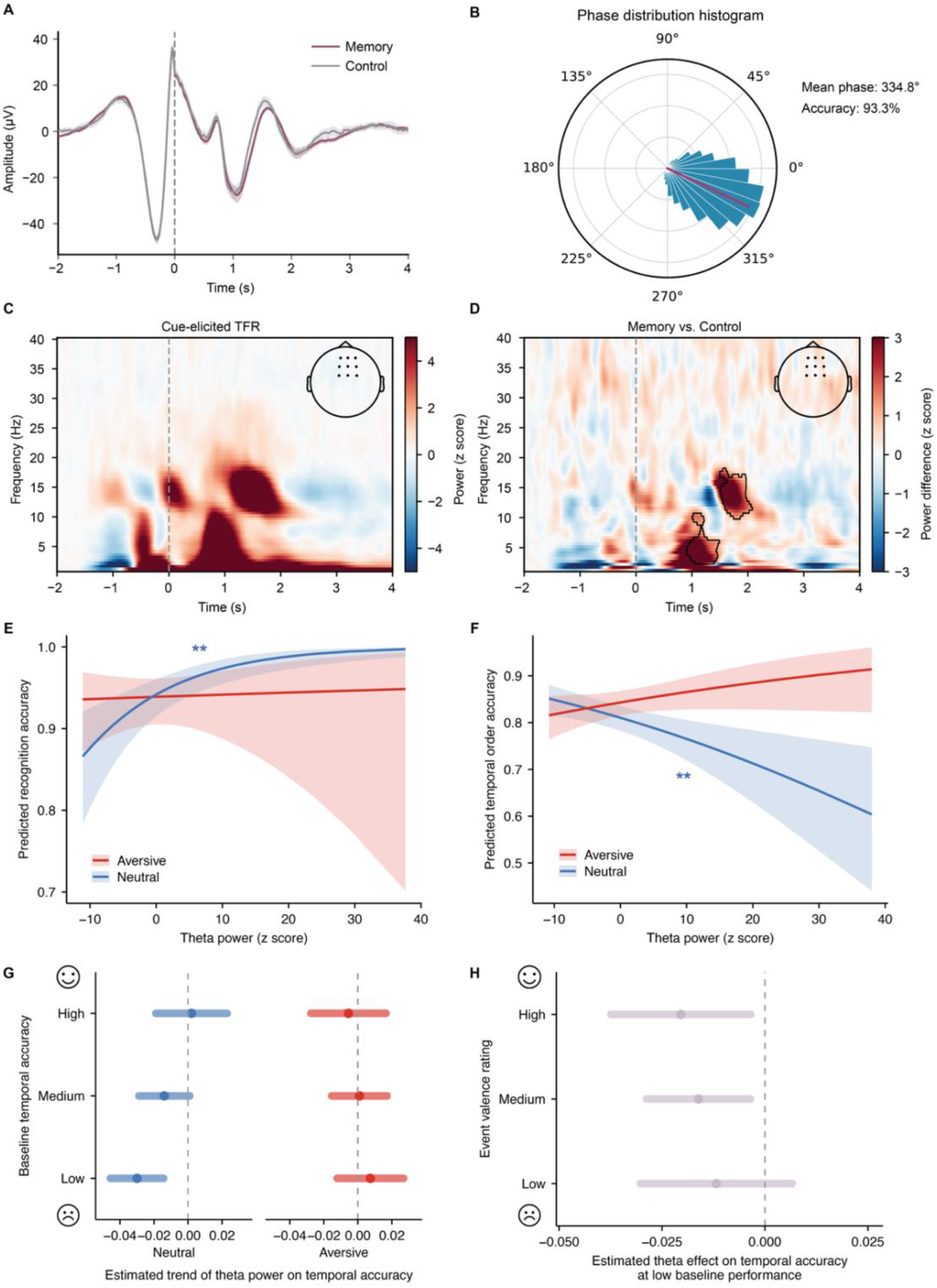
Cue-elicited neural activity during SWS and its associations with content and temporal memory. (A) Event-related potentials (ERPs) elicited by closed-loop auditory cues in the memory (aversive and neutral) and control conditions. (B) Phase distribution of closed-loop auditory cue delivery during sleep, with 93.3% of cues delivered at the SO up-state. (C) Cue-locked time–frequency power, collapsed across conditions, revealed prominent low-frequency activity before cue onset followed by increased sigma activity around cue delivery. (D) Time–frequency contrast between memory and control cues: memory cues elicited significantly greater frontocentral delta–theta and sigma power than control cues. (E) Greater cue-elicited theta power predicted higher recognition accuracy for neutral post-boundary items. (F) Cue-elicited theta power negatively predicted temporal-order accuracy for neutral events. (G) This theta–memory relationship was strongest when pre-sleep temporal memory was weak. (H) Greater theta power predicted poorer temporal-order memory for events rated as moderate to high in valence—that is, relatively less aversive. *p*_cluster_ < .05; \*\**p* < .01; Error bars represent 95% confidence intervals (CIs); CIs excluding zero indicate statistical significance at *p* < .05.

Unlike open-loop TMR procedures, the pre-cue baseline in our closed-loop design was characterized by prominent low-frequency activity followed by increased sigma activity around cue onset, consistent with the tendency of spindles to occur near the SO up-states (Fig. 3C; Ngo et al., 2013; Ngo & Staresina, 2022; Staresina, 2024). Following cue onset, both memory and control cues elicited increased low-frequency, sigma, and beta EEG power relative to baseline, indicating robust auditory-elicited neural responses during slow-wave sleep (*p*_clusters_ < .05, Supplementary Fig. S1). Critically, relative to control cues, memory cues elicited significantly greater delta-theta and sigma power over frontocentral electrodes (*p*_clusters_< .05; Fig. 3D), possibly indicating cue-triggered memory reactivation. In contrast, no significant differences were observed between aversive and neutral memory cues.

#### Cue-elicited theta power differentially tracked content and temporal memories

To understand the neural basis of the observed effects on content and temporal memory, we next examined how memory-specific EEG responses during sleep may be associated with post-sleep content and temporal memories. For each cue, the median power within each significant cluster was extracted per frequency band. Separate GLMMs were fit for the delta (0.5–4 Hz), theta (4–8 Hz), and sigma (11–16 Hz) bands, with EEG power, event valence (aversive vs. neutral), boundary position (pre- vs. post-boundary) and their interactions as fixed effects, while taking control-cue band power as a covariate with random intercepts for participant and event. For content memory, we found that memory-cue–elicited theta power showed a significant three-way interaction with valence and boundary position, *χ*²(1) = 3.93, *p* = .048, while control-cue theta power was not associated with recognition, *χ*²(1) = 0.12, *p* = .731. Simple-slope analyses revealed that greater memory-cue-elicited theta power was associated with higher recognition accuracy for neutral post-boundary items, *z* = 2.65, *p* = .008 (Fig. 3E), but not for neutral pre-boundary items, *z* = –0.05, *p* = .964, nor for aversive events at either boundary position (all *ps* ≥ .422). Direct slope comparisons further indicated that the theta–recognition association was stronger for neutral than for aversive post-boundary items, *z* = 2.04, *p* = .042, whereas the valence-specific slopes did not differ for pre-boundary items, *z* = –0.61, *p* = .544. A parallel pattern emerged for reaction times: greater theta power predicted faster responses for neutral post-boundary items, *z* = –2.30, *p* = .022, but not for their aversive counterparts, *z* = 0.67, *p* = .502. No comparable effects were observed in the delta or sigma bands (Supplementary Fig. S2A&C).

Using a parallel modeling approach with temporal-memory accuracy as the dependent variable, we added test time (immediate vs. delayed) as an additional fixed effect and included baseline performance and control-cue-elicited EEG responses as covariates. Results revealed that memory-cue–elicited theta power interacted with valence in predicting temporal memory, *χ*²(1) = 4.46, *p* = .035, while control-cue-elicited theta was not, *χ*²(1) = 1.26, *p* = .261. However, unlike recognition, here higher memory-cue-elicited theta power was associated with less accurate temporal memory for neutral events, *z* = –2.81, *p* = .005, whereas no reliable association emerged for aversive events, *z* = 0.27, *p* = .791 (Fig. 3F). No other theta-related interactions involving test time or boundary position were detected (all *ps* ≥ .247). Sigma band analyses showed a similar significant interaction between memory-cue-elicited sigma power and valence, *χ*²(1) = 4.83, *p* = .028: greater sigma power predicted poorer temporal-memory accuracy for neutral events, *z* = – 2.83, *p* = .005, but not for aversive events, *z* = 1.68, *p* = .092 (Supplementary Fig. S2F). No corresponding effects were observed in the delta band (Supplementary Fig. S2D). Results from models fitted separately by test time are provided in Supplementary Tables S3 (theta) and S4 (sigma).

Taken together, cue-elicited theta power showed opposing associations with content and temporal memory: stronger theta activity was positively associated with recognition of neutral post-boundary items, yet was negatively associated with temporal-order memory for neutral events. No reliable associations were observed for aversive memories in either memory domain. Notably, these effects were evident only for memory-cue-elicited EEG responses but not for control-cue-elicited EEGs, indicating that the observed associations were specific to memory reactivation rather than reflecting general auditory responses in sleep EEG.

### Non-preregistered exploratory analyses: Baseline temporal memory performance and valence rating qualify theta-memory associations

To examine whether baseline memory strength contributed to valence-dependent theta-memory association, we fitted an additional model that included pre-sleep baseline performance as a moderator. This analysis was motivated by the observation that aversive events were recalled with higher accuracy than neutral events at baseline (78.5% vs. 67.9%, *p* < .001), together with prior evidence that TMR effects depend on pre-sleep memory strength, with greater benefits typically observed for initially weaker memories (Creery et al., 2015; J. Liu et al., 2023; Schechtman et al., 2023b). Event boundary and test time were retained as control variables, as previous analyses revealed no theta-related interactions with these factors. This analysis revealed a significant three-way interaction among theta power, valence, and baseline temporal memory, *χ²*(1) = 6.76, *p* = .010. Follow-up analyses showed that, when baseline temporal memory was low (-1 SD), higher theta power predicted poorer temporal memory for neutral events, *z* = -3.75, *p* < .001, but not for aversive events, *z* = 0.74, *p* = .460; and the difference between the neutral and aversive slopes was significant, *z* = -3.03, *p* = .002 (Fig. 3G). The valence difference was not significant at mean or high baseline levels, all *ps* ≥ .161, indicating that the theta-memory relationship in neutral events emerged primarily when pre-sleep temporal memory was weak.

To further examine whether the neutral vs. aversive difference reflects finer-grained valence variation across events, we replaced the categorical valence factor (aversive vs. neutral) with event-level continuous pre-sleep ratings of valence and arousal in exploratory analyses. For arousal ratings, there was no evidence that arousal modulated the theta-memory relationship, as neither the theta × arousal interaction, *χ*²(1) = 0.30, *p* = .586, nor the theta × arousal rating × baseline interaction, *χ*²(1) = 2.07, *p* = .150, was significant. By contrast, for valence ratings, the theta × valence rating × baseline interaction was significant, *χ*²(1) = 5.10, *p* = .024, indicating that the association between cue-elicited theta power and later temporal memory depended on baseline temporal memory accuracy and was further modulated by valence ratings. Follow-up simple-slope analyses showed that when baseline temporal memory was low (−1 SD), higher theta power predicted poorer temporal memory for events at mean, *z* = -2.50, *p* = .012, and high valence *z* = - 2.36, *p* = .019 (i.e., less aversive), whereas the corresponding slope for low valence (i.e., more aversive) events was not reliable, *z* = -1.26, *p* = .209 (Fig. 3H). Moreover, sigma power showed a similar pattern, predicting poorer temporal memory for less aversive events (Supplementary Data S2), whereas no comparable effects were found for delta power. Together, these exploratory analyses suggest that the association between theta power and temporal-order memory depended on both baseline memory strength and subjective valence. Greater theta power was associated with poorer temporal-order memory primarily for events with lower baseline temporal memories and were perceived as more neutral or less aversive.

#### Non-preregistered analysis revealed event-specific neural representations during slow-wave sleep

Cue-elicited EEG power tracks the magnitude of neural responses during sleep but does not reveal whether these responses carry event-specific representations (Duan et al., 2025; J. Liu et al., 2023). We therefore employed multivariate representational similarity analysis (RSA) to characterize event-specific representations elicited by memory cues and to compare them with cue-specific representations elicited by control cues. Motivated by the theta–memory associations identified in the preceding power analyses, we focused on EEG data filtered in the theta band (4– 8 Hz). Event-specific representations were indexed by greater within-event (WE) than between-event (BE) similarity, with the three images associated with each pseudoword cue constituting a post-boundary subevent (Fig. 4A). Memory cues elicited significantly greater WE than BE similarity from 473 to 1,348 ms following cue onset (*p*_cluster_ < .001), whereas no significant clusters emerged for control cues (Fig. 4B). The direct contrast between memory and control cues, however, did not reach statistical significance (*p*_cluster_ > .05). Importantly, these results remained consistent after matching the number of trial pairs between WE and BE conditions and between memory and control cues (Supplementary Fig. S4). Together, these findings demonstrate distinct theta-band neural response patterns associated with individual memory cues during slow-wave sleep.

**Fig. 4.**
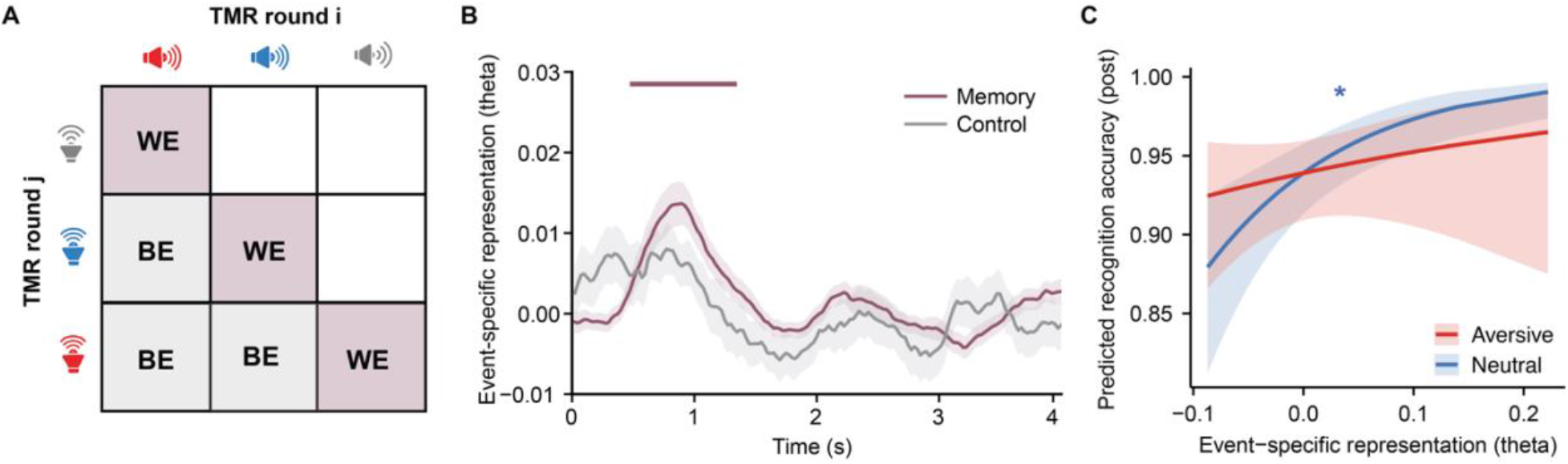
Event-specific neural representations following TMR cues during SWS. (A) RSA scheme across TMR blocks. Artifact-free theta band EEG patterns were correlated between trials sharing the same auditory cue (within-event similarity, WE) and between trials with different cues (between-event similarity, BE). Event-specific representations were quantified by contrasting WE and BE similarity. (B) RSA of theta-band EEG activity revealed significant event-specific representations in response to memory cues from 473 to 1,348 ms after cue onset. (C) Stronger event-specific representations predicted better recognition performance for neutral post-boundary items. *p*_cluster_ < .05; \**p* < .05.

#### Event-specific neural representations are associated with content memory for neutral post-boundary items

We next examined whether the strength of event-specific neural representations predicted subsequent content and temporal-order memory. For each memory cue, we extracted the median event-specific similarity value within the significant theta-band cluster (473–1,348 ms; Fig. 4B) and entered it into GLMMs paralleling those used in the preregistered EEG power analyses. Neither the main effect of representational strength nor its interactions reached significance in the omnibus model predicting content memory, all *ps* ≥ .179. Motivated by the previous results on theta-memory associations, we conducted exploratory focal analyses revealing that stronger neural representations were associated with better recognition for neutral post-boundary items, *z* = 2.10, *p* = .036, but not for their aversive counterparts, *z* = 0.58, *p* = .560 (Fig. 4C). By contrast, representational strength was not reliably associated with temporal-order memory accuracy (all *ps* ≥ .126). Together, these exploratory findings suggest that event-specific neural representations during slow-wave sleep selectively predicted subsequent content memory for cued items in neutral events, but were not reliably linked to temporal-order memory.

## Discussion

Memories of waking experiences are not exact replicas of the past. Rather, sleep selectively transforms different components of episodic memory, shaping both what is remembered and how events are reconstructed over time. In this preregistered study, we demonstrated that targeted memory reactivation during slow-wave sleep exerted distinct effects on memory for event content and for temporal structure. Reactivation induced faster and more confident recognition responses with more liberal recognition criteria. Notably, reactivation differentially modulated temporal-order memory based on time: it initially impaired, but subsequently preserved memories of event sequences. These behavioral effects were paralleled by sleep neurophysiology: for neutral events, stronger cue-elicited theta activity was associated with better recognition of event content, whereas greater theta activity predicted poorer temporal-order memory. No comparable associations were observed for aversive events. Nor did reactivation reliably alter affective responses to the reactivated events. Together, these findings challenge the view that sleep reactivation uniformly strengthens episodic memories. Rather, sleep reactivation selectively prioritizes the consolidation of event content while simultaneously reshaping the consolidation trajectory of temporal memories.

We first documented the cueing benefits, evidenced by faster, more confident, and more liberal response biases in recognition, aligning with mounting evidence that TMR during non-rapid eye movement (NREM) or slow-wave sleep consolidates declarative memories (Carbone & Diekelmann, 2024; Hu et al., 2020). At the mechanistic level, rodent work indicates that external cues can bias the content of hippocampal replay (Bendor & Wilson, 2012), possibly via cue-elicited cortical activity during NREM sleep that shapes which memories—and potentially which sequences—are selected for subsequent hippocampal reactivation (Lewis & Bendor, 2019; Rothschild et al., 2017). Human studies have likewise shown that TMR can elicit detectable memory reactivation during sleep (Cairney et al., 2018; Schreiner et al., 2018; Schreiner & Staudigl, 2020; B. Wang et al., 2019), thereby supporting subsequent memory consolidation and providing a means of experimentally probing this process. Our findings extend this approach to complex event memories, showing that closed-loop cueing at SO up-states also facilitated access to event content one week later. Notably, we observed comparable cueing benefits on recognition for both pre- and post-boundary items, suggesting that cueing benefits generalized to uncued pre-boundary items within the same cued event. This pattern is compatible with recent evidence indicating that reactivation would simultaneously benefit multiple contextually linked memories (Schechtman et al., 2023b). Intriguingly, stronger memory-cue-elicited theta power and event-specific neural representations were associated with higher recognition only for neutral post-boundary items, suggesting that cue-elicited theta specifically tracked reactivation of directly cued memories even when cueing benefits would generalize broadly across the whole event.

Instead of being isolated episodes, memories are temporally orchestrated. Contrary to our prediction that reactivation would strengthen temporal memories, we found that cueing initially impaired temporal-order memories. This impairment may reflect a transient disruption of the temporal scaffold in which the reactivated content was embedded. Although our design did not allow us to determine the sequential order of replay, the observed impairment suggests that multiple associated items within an event can be reactivated without faithfully reinstating their original sequence (Schechtman et al., 2021). At the neural level, EEG–behavior analyses further revealed a striking dissociation for neutral events: stronger cue-elicited theta activity predicted better recognition of cued post-boundary items, yet poorer temporal-order memory for the same events, with the disruptive effect on temporal order emerging most prominently at the early test. Thus, rather than faithfully reinstating the original temporal structure, TMR may preferentially reactivate multiple event elements in parallel, thereby blurring their sequential order. More broadly, these findings indicate that exogenous reactivation can selectively reshape specific memory features rather than uniformly strengthening the entire event—for instance, enhancing distinctive features while weakening shared ones to promote differentiation between overlapping representations (Siefert et al., 2024).

However, this cue-induced disruption was transient: cueing attenuated the decline in temporal-order memories over the subsequent week. This reversal suggests that the initial perturbation of temporal structure may have been followed by further consolidation on longer timescales. Specifically, event memories tagged by TMR may undergo additional endogenous processing across subsequent periods of sleep and wakefulness, potentially restoring and stabilizing their temporal relationships within the broader event structure. This possibility is consistent with evidence that post-encoding sleep durably preserves temporal-order memory, with effects persisting even up to one year later (Diamond et al., 2025; Drosopoulos et al., 2007; Griessenberger et al., 2012). Moreover, sleep-related improvements in temporal-order memory have further been linked to SO–spindle coupling, a consolidation mechanism implicated in memory stabilization and integration (Cross et al., 2025; Diamond et al., 2025; Staresina, 2024). Together, these findings point to a two-stage “reorganization–consolidation” account: TMR may transiently disrupt the temporal embedding of reactivated memories while simultaneously tagging them for subsequent endogenous reactivation, which in turn preferentially consolidates the temporal structure of the tagged sequences.

Examining the neural activity underlying cue-triggered memory reactivation further revealed a valence-dependent neural-behavioral association. Although valence (i.e., aversive vs. neutral) did not moderate the behavioural effects of TMR, cue-elicited theta activity showed the clearest negative associations with both content and temporal memory for neutral but not aversive events. Our exploratory follow-up analyses further indicated that the stronger cue-elicited theta was associated with poorer temporal memory, which was most pronounced when pre-sleep temporal memory was weak and when events were rated as less aversive—an effect better captured by subjective valence than by arousal ratings. Thus, this result stands in contrast with previous research revealing that sleep reactivation would preferentially benefit weakly encoded memory contents (Creery et al., 2015; J. Liu et al., 2023; Petzka et al., 2021; Schapiro et al., 2018; Tambini et al., 2017), again supporting the notion that memory content and temporal memories can be differentially influenced by sleep reactivation. In the present study, aversive events were better remembered before sleep, possibly because emotional salience prioritizes memory formation through coordinated amygdala–hippocampal encoding processes (Costa et al., 2022; Qasim et al., 2023; H. Zhang et al., 2024). Their more stable pre-sleep representations may therefore have rendered aversive memories less amenable to further modification by exogenously triggered reactivation. By contrast, neutral events—particularly those with weakly encoded temporal relations—may have remained more labile and thus more open to reactivation-induced reorganization. Together, these findings suggest that the consequences of sleep reactivation depend not only on emotional valence per se but also on the stability of pre-sleep memory traces, with the greatest modification occurring for relatively weak, less aversive memories.

Contrary to our preregistered prediction that sleep reactivation would attenuate affective responses, we observed no reliable changes in either subjective valence or arousal ratings. This null finding aligns with a broader pattern in the NREM TMR literature: cueing consistently modulates the consolidation of emotional memories (Cairney et al., 2014; Lehmann et al., 2016; Yuksel et al., 2025) or alters fear-extinction learning (Hauner et al., 2013; Zheng et al., 2024) rather than changing subjective valence or arousal (Hu et al., 2020). By contrast, emerging research shows that cueing emotional memories during REM sleep reduces subjective arousal while attenuating neural markers implicating emotional reactivity (Greco et al., 2025; Hutchison et al., 2021; Patriota et al., 2026), suggesting that the affective consequences of memory reactivation may depend on different sleep stages (i.e., NREM vs. REM) (Cabrera et al., 2024; Goldstein & Walker, 2014). Moreover, when NREM reactivation does attenuate affective responses, these benefits appear to depend on reactivating competing positive memories rather than on reinstating the original aversive content (Recher et al., 2024; Xia et al., 2024). Our null finding suggests that reactivating aversive event memories during NREM sleep may preferentially consolidate the declarative component of emotional experiences without necessarily altering their affective tone.

Limitations and open questions warrant discussion. First, identifying sequential memory reactivation during sleep in humans remains challenging (Y. Liu et al., 2022; Schreiner & Staudigl, 2020). Our mechanistic interpretation—particularly for temporal-order memory—therefore remains indirect. Future studies could combine TMR with high-resolution recordings such as intracranial EEG and sequence-sensitive multivariate decoding methods to delineate hippocampal replay of event sequences and how such replay shapes the temporal scaffolding of event memories. Second, although the time-dependent effect on temporal-order memory is consistent with a “reorganization–consolidation” account, direct evidence for this two-stage process is still lacking. Future studies could track memory transformation over time, examining how initially disrupted temporal structures are subsequently consolidated into more coherent event representations.

In conclusion, sleep reactivation does not simply preserve event memories as they were encoded. This preregistered study revealed that although reactivation facilitated recognition, it dynamically reshaped how people remembered the temporal structure in which memory was embedded. Moreover, cue-elicited theta activity emerged as a neural signature tracking these behavioral effects, pinpointing a candidate mechanism for event-memory reactivation during sleep. Together, these behavioral and neurocognitive evidences highlight a selective, time-dependent sleep reactivation mechanism that reorganizes temporally structured event memories.

## Methods

### Participants

Thirty-five participants (mean age = 23.01 ± 1.75 years; 29 female) recruited from the University of Hong Kong were included in the final analyses. Data from 24 additional participants were excluded due to: inability to reach 80% accuracy during learning (n = 2), or completion of fewer than seven rounds of auditory cue administration during SWS (n = 20), and technical problems (n = 2). Four participants did not attend the delayed test session, resulting in thirty-one participants for the delayed test analyses. Participants met the following eligibility criteria: age 18-25 years, no chronic or acute medical conditions, and good sleep quality. Qualified participants were asked to maintain regular sleep schedules before and during the experiment. The sample size of the current study was comparable to, or larger than, those reported in previous human closed-loop sleep TMR studies (*N* = 11 to 34, Navarrete et al., 2024; Ngo et al., 2013; Ngo & Staresina; Santamaria et al., 2024; Siefert et al., 2024). The study was approved by the Human Research Ethics Committee of the University of Hong Kong.

### Materials

The study included 16 aversive events and 16 thematically matched neutral events, each composed of six images extracted from 2-3minute film clips to ensure a coherent narrative context (Y. Zhang et al., 2026). Within each event, each image depicted relatively distinct scenes that were not directly connected, but together they formed a meaningful narrative. In aversive events, the first three images portrayed neutral contents, followed by three images depicting aversive scenes. In contrast, neutral sequences consisted of six images depicting difference neutral content. This design enabled an emotional shift within aversive sequences, allowing investigation of the retroactive impact of emotional subevents on memory for preceding neutral subevents. To pair with these subevents, sixty-eight two-syllable pseudowords were created by randomly pairing two neutral Chinese characters (Xia et al., 2022). Vocalizations of the pseudowords were generated in Mandarin using the Microsoft Azure Text-to-Speech function (word duration: *M* = 1.226 s, *SD* = 0.334 s). Sixty-four pseudowords were randomly paired with one subevent (i.e., images 1-3 or images 4-6 within each event) to serve as memory cues, while the remaining four served as control cues. Manipulation checks using the emotional compass ratings (see learning task below for details) confirmed that aversive events were rated as significantly more negative (*M* = -0.094, *SD* = 0.037) and arousing (*M* = 0.040, *SD* = 0.087) than neutral events (valence: *M* = 0.040, *SD* = 0.046; arousal: *M* = -0.085, *SD* = 0.075), *ts*(34) > 15.36, *ps* < .001.

### Experimental design and procedure

The experiment consisted of an overnight TMR session and a one-week delayed session. In the TMR session, participants arrived at the sleep laboratory around 8:00 PM and were first prepared for EEG recordings. Afterward, participants completed a series of tasks, including psychomotor vigilance task, cue familiarization, event learning and an immediate retrieval test. They then went to bed, and the lights were turned off at midnight. TMR commenced after 5 minutes of stable SWS. After approximately 8 hours of sleep (around 8:00 AM), participants were awakened. Thirty minutes after awakening (to mitigate sleep inertia), participants completed the post-sleep memory test, which was identical to the pre-sleep test. One week later, participants returned to the laboratory for a delayed memory test, which included an old/new recognition task in addition to the previously administered tasks.

### Experimental tasks

#### Psychomotor vigilance task (PVT)

Participants engaged in a 5-minute PVT before and after sleep to evaluate their vigilance. They were instructed to focus on a fixation cross on the screen, which switched to a counter counting upward from 0s. Participants were required to stop the counter as quickly as possible by pressing the space bar. The counter continued until a response was registered or until it reached a maximum of 2s. The inter-trial interval (ITI) varied randomly between 2 and 10s. After each trial, feedback displaying the reaction time was provided.

#### Cue familiarization task

Prior to the encoding session, participants completed a cue familiarization task in which all memory cues were played once to facilitate subsequent learning of the pseudoword–image associations. Each trial began with a fixation being presented for about 1s, followed by the simultaneous auditory and visual presentation of a pseudoword cue. Participants shall press a key to indicate whether they had heard the cue clearly or not ("yes" or "no"). This procedure ensured perceptual clarity and attentional engagement before proceeding to the learning phase.

#### Event learning task

The learning phase comprised two to four encoding–recognition cycles. All participants completed at least two cycles, with additional cycles (up to a maximum of four) administered as needed until recognition accuracy reached 80%. This criterion ensured sufficient learning strength for the subsequent experimental manipulations. In each cycle, participants encoded 32 events (16 aversive, 16 neutral), with each event comprising six images presented sequentially for 2s each, accompanied by pseudowords serving as memory cues. The first three and last three images in each event are associated with each of the two pseudowords, with the second cue used for later TMR cueing. This divided each sequence into two subevents, with the second cue also serving as an event boundary. Participants were instructed to form meaningful associations and mentally construct a coherent narrative linking all six images. In the first cycle only, immediately after viewing each sequence, participants rated the event using an emotional compass—a continuous two-dimensional affective space with valence (negative to positive) on the horizontal axis and arousal (low to high) on the vertical axis (McClay et al., 2023). Participants next recalled each image in their mind and rated emotional feelings toward each recalled image by moving the mouse continuously within the compass area to indicate how they remembered feeling. The entire mouse trajectory was tracked, and the cursor coordinates at each click were recorded, capturing the temporal dynamics of emotional responses throughout the event. Additionally, participants rated the coherence of their self-constructed narrative on a 9-point Likert scale.

Following the encoding phase in each cycle, participants completed a forced-choice recognition test consisting of 64 trials (16 events × 2 valence [aversive, neutral] × 2 boundary [pre, post]). In each trial, a randomly selected memory cue was presented both aurally and visually, and the target triplet (the subevent originally paired with that cue) was presented alongside three foil triplets. Participants were instructed to select the correct triplet. Foils were randomly selected but matched to the target in emotional valence and boundary position, ensuring that all options were perceptually and contextually similar. This design increased task difficulty and minimized reliance on categorical features. Regardless of the accuracy, correct feedback was provided after each trial to support associative learning. The second, third, and fourth cycles followed the same procedure but omitted the emotional compass ratings, focusing solely on reinforcing narrative construction and pseudoword-image associations through repeated encoding-recognition cycles.

#### Pre-/Post-TMR memory test

To evaluate memory changes across sleep and TMR, participants completed two tasks before and after the sleep session: 1) a cue rating task, and 2) a temporal-order memory task. In the cue rating task, all 64 pseudoword cues were tested. On each trial, a pseudoword cue was presented, and participants were instructed to mentally replay the associated sub-events within 3s. They then made a single click on the emotional compass to indicate their emotional response to the retrieved memory. Pre- and post-sleep differences in emotional ratings were analyzed to assess changes in memory-specific emotional responses, with particular attention to how cueing influences these changes. In the temporal memory task, all possible image pairs from each event were presented sequentially. Specifically, these included three pre-boundary image pairings (i.e., 1-2, 1-3, 2-3), three post-boundary image pairings (i.e., 4-5, 4-6, 5-6), and three cross-boundary image pairings (i.e., 1-4, 2-5, 3-6). For each pair, participants first indicated which image occurred earlier in the event, followed by a rating of the subjective temporal distance between the two images on a 0-4 scale (0 = “very close,” 4 = “very far”). Changes in performance from pre- to post-sleep were analyzed to evaluate how TMR affected the temporal-order memory.

#### Delayed memory test

Seven days after the initial session, participants returned to the laboratory to complete a delayed memory test consisting of four tasks: a cue valence/temporal judgment task, a cue rating task, an old/new recognition task, and a temporal-order memory task. This session was designed to evaluate sleep and TMR’s long-term impact on episodic memories. In the old/new recognition task, participants were presented with a total of 384 images, comprising 192 previously encoded images (16 sequences × 2 valence [aversive, neutral] × 6 images) and 192 novel foils.

Each trial began with a 1.5s fixation cross, followed by the display of a single image. Participants were instructed to indicate as quickly and accurately as possible whether the image was “old” (previously studied) or “new” (unseen). Following this binary judgment, participants provided a confidence rating on a 5-point scale ranging from 0 (guessing) to 4 (very certain). The judgment, cue rating, and temporal-order memory tasks followed the same procedures as in the pre-/post-TMR session, allowing for within-subject comparisons of memory performance across immediate and delayed retrieval intervals.

#### Closed-loop TMR during SWS

The closed-loop TMR procedure was conducted during an overnight sleep session starting at approximately 12:00 a.m. Participants slept in a laboratory bedroom for up to 8 hours while their brain activity was continuously monitored using high-density electroencephalography (EEG). Throughout the night, pink noise was played as background sound to mask external disturbances and maintain acoustic consistency. TMR was implemented in a closed-loop design, with auditory cues precisely time-locked to the UP state of slow oscillations (SO) detected during slow-wave sleep (Ngo & Staresina, 2022). Real-time EEG monitoring and closed-loop stimulation were carried out using OpenViBE platform (Renard et al., 2010). Sleep staging was performed based on EEG signals recorded at Fz, Cz, and Pz, along with EOG and EMG, referenced to M1 and M2. Trained experimenters continuously monitored the EEG to identify stable SWS sleep and trigger cue delivery accordingly.

To identify the SO up-state, the filtered signal (0.5-4 Hz) from electrode Fz had to rise, cross a threshold of +35 μV, and show a change in slope from positive to negative, indicating a local maximum. Each time an UP state was detected, a cue was delivered through a loudspeaker located within the room, and the detection protocol paused for 6 s. The sound pressure level was maintained at 39-40 dB SPL, measured at the position of the participant’s pillow. During the TMR procedure, 16 previously learned pseudoword cues (8 aversive, 8 neutral) were presented along with 4 novel pseudowords serving as non-memory control cues. Each block consisted of these 20 cues delivered in random order, followed by a 30 s pause before the next block began. Cueing commenced after at least 5 minutes of stable SWS had been observed. If micro-arousals, awakenings, or transitions to N1 or REM sleep were detected, cueing was immediately paused and resumed only when stable SWS was observed. The cueing session continued until either 20 rounds were completed or till 3:00 a.m. Participants who received fewer than 7 rounds were excluded from analysis.

To assess the precision of the closed-loop algorithm in targeting the SO up-state, raw EEG data were processed with a zero-phase band-pass filter (0.1-40 Hz), and the SO phase was estimated from the 0.5-4 Hz band using the Hilbert transform. A cue was classified as successfully delivered during the SO up-state if its onset phase fell within ±90° of the positive SO peak (i.e., 270°-90°); all other cues were classified as falling outside the up-state. Up-state targeting accuracy was computed for each participant as the proportion of valid cues falling within this interval and was then averaged across participants. The closed-loop algorithm exhibited high phase-targeting precision: the pooled cue-onset phase distribution yielded a circular mean of 334.8° (bootstrap 95% CI: 332.7°-337.5°), and, on average, 93.3% of valid cues per participant were delivered within the predefined SO up-state window.

#### EEG Recording

EEG was recorded using a 64-channel EEG cap connected to an eego amplifier (ANT Neuro), with electrodes arranged according to the international 10-20 system and referenced online to CPz. Two electrooculogram (EOG) electrodes were placed diagonally around the eyes— one below the left eye and one above the right eye—to monitor horizontal and vertical eye movements. Additionally, two bipolar electromyogram (EMG) electrodes were positioned on the chin to record muscle activity. EEG signals were sampled at 500 Hz throughout both wakefulness and sleep. During wakefulness, the EEG signal was band-pass filtered online between 0.3 and 30 Hz for monitoring purposes. During sleep, EEG data from the eego system were streamed to the OpenViBE Brain Computer Interface software for continuous recording and closed-loop TMR manipulation.

## Data analysis

### EEG data processing

EEG data were processed using the MNE-Python toolbox (Gramfort et al., 2013). Preprocessing steps included down-sampling to 200 Hz, application of a 50 Hz notch filter, and bandpass filtering between 0.1 and 40 Hz using a finite impulse response (FIR) filter. Noisy or malfunctioning channels were visually identified and marked, followed by re-referencing to the average of M1 and M2. Continuous EEG recordings were segmented into epochs from -15s to +15s relative to cue onset for subsequent time-frequency analysis and event detection. Artifactual epochs were excluded based on visual inspection, and marked bad channels were interpolated before further analysis.

### Sleep staging analysis

Sleep staging was performed using the Yet Another Spindle Algorithm (YASA), a machine learning-based method implemented in Python (Vallat & Walker, 2021). Raw overnight EEG data were re-referenced to the average of [M1, M2], following YASA’s recommended configuration. The algorithm was applied using data from the C4 electrode (or C3 if C4 was identified as a bad channel), along with EOG and EMG channels. The automatically generated hypnograms were checked by a trained experimenter (Supplementary Table S5).

### Event-related potentials (ERPs)

To assess the brain response elicited by memory and control sounds, we calculated ERPs by segmenting the 0.1- to 40-Hz band pass-filtered EEG signal into 6 s epochs time locked to the cue onset (i.e., -2 to +4 s relative to cue onset). ERPs were baseline-corrected using the pre-cue interval from -2 to -1.5 s, a relatively stable period before SO UP state, and were then averaged across trials for each condition.

### Time-frequency analysis (TFR)

For time-frequency analysis, EEG epochs were cropped to the interval from -3 to +5 s relative to TMR cue onset. A continuous wavelet transformation featuring variable cycles (3 cycles in length at 1 Hz, increasing linearly with frequency up to 15 cycles at 40 Hz) was applied to the cropped epochs. To reduce edge artifacts, the resulting time-frequency data were further trimmed to -2 to +4 s relative to cue onset. Spectral power was Z-score normalized across trials within each condition using a baseline window from -2 to -1.5 s. Trial-level time-frequency representations were retained for subsequent analysis. We then performed statistical tests on the median power within the nine pre-defined frontocentral channels (F1/2, FC1/2, C1/C2, Fz, FCz and Cz) in each condition (e.g., aversive/neutral memory cues vs. control sounds). These nine channels were selected per previous studies examining auditory/memory processing during sleep (Chen et al., 2024; Xia et al., 2022; Yao et al., 2024; Züst et al., 2019). To assess cue-related neural responses, memory cues were contrasted either with pre-cue baseline activity or with control sounds. A two-dimensional temporal-frequency permutation test (two-tailed, one-sample with randomization of 1000 and a statistical threshold of 0.05) was utilized to evaluate significant clusters.

### Representational similarity analysis (RSA)

To quantify the cue-elicited neural representations during sleep, we performed RSA on the raw EEG activity between every 2 clean TMR trials that were drawn from different TMR rounds to minimize correlations due to the temporal proximity. Motivated by the theta–memory associations identified in the preceding power analyses, RSA was performed on EEG data band-pass filtered in the theta band (4–8 Hz). To characterize the temporal dynamics of neural representations, RSA was applied using sliding time windows of 500 ms with a step size of 25 ms, yielding feature vectors of 301 × 58 (time points × channels) for each time window. For each window, representational similarity was computed as the Spearman correlation between the vectorized EEG features of every pair of trials from different TMR rounds. All correlation coefficients were Fisher *z*-transformed prior to statistical analysis.

To quantify event-specific representations, we calculated within-event (WE) similarities based on the neural similarities of whole-brain EEG activity patterns between the same cues, and between-event (BE) similarities based on the neural similarities of whole-brain EEG activity patterns between different cues. Event-specific neural representations were quantified as the difference between within-event and between-event similarities (WE – BE). Per participant, we obtained 869 ± 464 WE and 6,080 ± 3,251 BE trial pairs for aversive cues, 880 ± 472 WE and 6,159 ± 3,303 BE trial pairs for neutral cues, and 437 ± 234 WE and 1,310 ± 701 BE trial pairs for control cues. WE and BE similarity values were first averaged across trial pairs within each participant, and event-specific representations were then assessed by contrasting the mean WE and BE values (i.e., WE – BE). For memory cues, event-specific representations were computed by averaging the WE – BE values of the aversive and neutral conditions. For statistical analyses, the WE – BE time courses were tested against zero across participants for memory and control cues, respectively. Multiple comparisons across consecutive time windows were corrected using cluster-based nonparametric permutation tests (one-tailed, one-sample with randomization of 1000 and a statistical threshold of 0.05).

To rule out the possibility that event-specific representations were driven by unequal trial pairs, we performed 2 control analyses. In the first analysis, we balanced the WE and BE trial pairs within each condition by randomly selecting an equal number of BE trial pairs to match the number of WE trial pairs. In the second analysis, we applied a stricter joint-balancing procedure in which trial pairs were matched across the memory and control conditions, while also ensuring equal numbers of WE and BE trial pairs. The significant time windows remained consistent across both control analyses, suggesting that the observed event-specific representations were not explained by unbalanced trial pairs (Supplementary Fig. S4).

### Statistics

To examine the immediate and delayed effects of TMR on emotional responses and temporal-order memory, we fitted mixed-effects models to trial-level data. Valence and arousal ratings were analyzed using linear mixed-effects models (LMMs), whereas binary temporal-order responses (correct vs. incorrect) were analyzed using binomial generalized linear mixed-effects models (GLMMs). Fixed effects included TMR status (cued vs. uncued), emotional valence (aversive vs. neutral), boundary condition (pre-, post-, or cross-boundary), test time (immediate vs. delayed), and all interactions. The corresponding pre-sleep baseline measure was included as a covariate. All models included random intercepts for participants and events to account for repeated observations. The models were implemented using the lmer() and glmer() functions from the lme4 package (version 1.1.33) in R (version 4.2.1). The general model formula was:

Valence rating/Arousal rating/Temporal order choice ∼ 1 + TMR × valence × boundary × time + baseline choice + (1 | participant) + (1 | event). Follow-up pairwise contrasts and simple-effects analyses were conducted using the *emmeans* package.

For long-term event recognition measures derived from signal detection theory, we conducted repeated-measures ANOVAs with TMR (cued vs. uncued), valence (aversive vs. neutral), and boundary (pre- vs. post-boundary) as within-subject factors. Because signal detection measures were derived from hit and false-alarm rates within each participant and experimental condition, analyses of recognition sensitivity (d′) and response criterion were conducted using participant-by-condition estimates. ANOVAs were implemented in R using the afex package. Reaction times and confidence ratings were analyzed using the same repeated-measures ANOVA framework.

To link cue-elicited neural responses during sleep to subsequent memory performance, we averaged EEG activity across repetitions of each cue to obtain subevent time-frequency representations and representational similarity measures. We then extracted median spectral power or representational strength from the significant time-frequency or RSA clusters identified in prior analyses. Because cue-elicited neural predictors were defined at the subevent level, subsequent memory outcomes were modeled at the same level. Performance was therefore expressed as the number of correct versus incorrect responses associated with each subevent and entered as a two-column matrix of successes and failures in binomial GLMMs. Fixed effects included EEG power or representational strength, valence, boundary, and their interactions; test time was additionally included for temporal memory. Baseline memory performance and control cue-elicited neural responses were included as covariates. All models included random intercepts for participant and event. All EEG predictors and continuous covariates were mean-centered before model fitting. The general model formulas were as follows:

Recognition choices ∼ EEG power/representation × valence × boundary + baseline choices + control power/representation + (1 | participant) + (1 | event).

Temporal order choices ∼ EEG power/representation × valence × boundary × time + baseline choices + control power/representation + (1 | participant) + (1 | event);

Temporal order choices ∼ EEG power × valence × baseline choices + boundary + time + control power + (1 | participant) + (1 | event);

Temporal order choices ∼ EEG power × valence/arousal rating× baseline choices + boundary + time + control power + (1 | participant) + (1 | event);

Statistical significance was evaluated at α = .05, and follow-up simple-effects analyses were used to interpret significant interactions.

## Supporting information

This document contains supplementary tables (1-5), figures (1-4), and notes (1-4) accompanying the main text.

## Acknowledgment

We thank Yingqian Chen (Christine) for her assistance in the data collection. The research was supported by the Ministry of Science and Technology of China STI2030-Major Projects (No. 2022ZD0214100), National Natural Science Foundation of China (No. 32171056), General Research Fund (No. 17614922) of Hong Kong Research Grants Council, and The University of Hong Kong University Research Committee Internal Seed Fund for Basic Research to X. H.

