## Supplementary material for "Sleep reactivation shapes memories of event and its temporal structure": This document contains supplementary tables (1-5), figures (1-4), and notes (1-4) accompanying the main text.

**This PDF file includes:**

Supporting methods and data
Figures S1 to S4
Tables S1 to S5

**Supplementary Methods**

Table S1. Preregistered main hypotheses and corresponding evidence from the current data.

| Preregistered hypothesis | Evidence from the current data |
|---|---|
| H1: TMR would strengthen memories of cued event content | Behavioral: TMR enhanced memory for cued event content, as reflected by faster responses, higher confidence, and a more liberal tendency to endorse "old" judgments.<br><br>Physiological: TMR-related memory benefits were more strongly predicted by theta activity for neutral cued subevents. |
| H2: TMR would consolidate temporal memory for neutral events but impair temporal memory for aversive events. | Behavioral: TMR produced short-term disruption but longer-term preservation of temporal memory.<br><br>Physiological: Stronger theta activity predicted poorer temporal memory for neutral events, whereas no such association was observed for aversive events. The theta–temporal memory relationship emerged most prominently for initially weak and subjectively less aversive events. |
| H3: TMR would attenuate emotional responses to cued aversive subevents. | TMR did not reliably modulate emotional ratings. Instead, ratings were primarily shaped by the designed event valence and boundary structure, with the strongest effects observed for aversive post-boundary subevents (Supplementary Data S1). |

Table S2. Deviations from preregistered analyses.

| Analysis | Deviation from preregistered analyses and rationale |
|---|---|
| Behavioral analysis | The preregistration specified both participant-level ANOVAs and trial-level mixed-effects models. In the final report, trial-level models served as the primary analyses, except for the recognition task, which was analyzed using signal detection measures. Mixed-effects models provide a more sensitive and appropriate test of behavioral performance by accounting for trial-level variability and random effects of participants and items. Pre-sleep performance was additionally included as a covariate in the GLMMs/LMMs to control for baseline differences between aversive and neutral events. |
| Time-frequency analysis | Given the closed-loop TMR design, we used –2 to –1.5 s as the baseline window to obtain a cleaner pre-stimulus estimate that was less likely to be contaminated by endogenous slow-oscillation (SO) activity. Formal analyses focused on predefined ROI channels based on prior TMR work, while exploratory three-dimensional cluster-based analyses revealed convergent theta-band effects. |
| Multivariate analysis | Instead of category-level decoding, we employed representational similarity analysis (RSA) to quantify event-specific neural representations. RSA was better suited to the present question, as it provides a continuous, trial-pair-based measure of event-specific representation that can be directly related to subsequent behavioral outcomes. |
| EEG-behavioral analysis | Mixed-effects models were selected according to the distributional properties of each dependent variable, with LMMs for continuous outcomes and GLMMs for non-Gaussian outcomes such as item-level accuracy.<br><br>Additional analyses examined whether TMR effects depended on baseline performance and whether subjective valence/arousal ratings captured affective modulation better than categorical valence. For item-level EEG–behavior analyses, EEG predictors (including cue-elicited power and representational strength) were summarized using median values to obtain more stable estimates. |

### Supplementary Results

#### Data S1. TMR did not modulate event-related emotional responses

We fit item-level linear mixed-effects models predicting cue emotional ratings from TMR status (cued vs. uncued), event valence (aversive vs. neutral), boundary position (pre-boundary vs. post-boundary), and test time (immediate vs. delayed), including all interactions. Pre-sleep baseline ratings were included as a covariate, with random intercepts for participant and event. For both valence and arousal ratings, TMR showed neither a main effect nor any interactions (all *ps* > .340), indicating that cueing did not reliably alter event-related emotional responses.

For valence ratings, a significant three-way interaction among test time, event valence, and boundary position emerged, $F(1, 4396.7) = 20.48$, $p < .001$. Across both tests, valence differentiation was confined to the the post-boundary portions of events, although this pattern attenuated at the delayed test. Specifically, aversive events were rated as more negative than neutral events at the post-boundary position at both tests (immediate: $z = 14.91$, $p < .001$; delayed: $z = 7.96$, $p < .001$), whereas no such difference emerged at the pre-boundary position (all *ps* > .10). Consistent with this pattern, within aversive events, post-boundary items were rated as more negative than pre-boundary items at both tests (immediate: $z = 14.64$, $p < .001$; delayed: $z = 6.55$, $p < .001$), whereas no boundary difference was observed for neutral events (all *ps* > .55).

For arousal ratings, a significant test time × event valence × boundary position interaction was also observed, $F(1, 4395.5) = 6.28$, $p = .012$. At the immediate test, aversive events elicited greater arousal than neutral events at both the pre-boundary ($z = -2.53$, $p = .011$) and post-boundary positions ($z = -15.56$, $p < .001$). By the delayed test, however, this difference persisted only at the post-boundary position ($z = -7.89$, $p < .001$) and was no longer detectable at the pre-boundary position ($z = 0.69$, $p = .489$). Within aversive events, post-boundary items elicited greater arousal than pre-boundary items at both tests (immediate: $z = -15.52$, $p < .001$; delayed: $z = -7.53$, $p < .001$). Overall, TMR did not modulate emotional responses. Instead, ratings were primarily shaped by designed event valence and boundary, with the strongest effects for post-boundary portions of aversive event.

**Data S2. Baseline temporal memory performance and valence rating qualify sigma-memory associations.**

To examine whether the relationship between cue-elicited sigma power and subsequent temporal memory was further moderated by baseline temporal memory, as observed for theta power, we fit a binomial GLMM including cue-elicited sigma power, valence, baseline temporal memory, and their interactions as predictors, with event boundary, test time and control sigma response included as control variables. This analysis revealed a significant interaction between sigma power and valence, $\chi^2(1) = 8.65$, $p = .003$, indicating that the relationship between cue-elicited sigma activity and later temporal memory differed between neutral and aversive events. The three-way interaction among sigma power, valence, and baseline temporal memory was marginal, $\chi^2(1) = 3.84$, $p = .050$, providing suggestive evidence that this valence-dependent relationship also varied with baseline performance. Follow-up simple-slope analyses showed that, at low baseline levels (-1 SD), greater sigma power predicted poorer temporal memory for neutral events, $z = -3.63$, $p < .001$, but better temporal memory for aversive events, $z = 1.87$, $p = .062$. The slopes differed significantly between neutral and aversive events at this low-baseline level, $z = -3.92$, $p < .001$. Similar valence differences in the sigma-memory slope were observed at mean baseline levels, $z = -2.94$, $p = .003$, but not high baseline levels (+1 SD), $z = -0.95$, $p = .342$ (Fig. S3A). Thus, cue-elicited sigma power showed a valence-dependent association with subsequent temporal memory, with the observed pattern appearing strongest for events that were initially remembered less accurately.

To further examine whether this valence-dependent pattern reflected more fine-grained emotional variation across events, we replaced the categorical valence factor with continuous valence and arousal ratings for each event in exploratory analyses. For arousal, there was no evidence that arousal reliably modulated the sigma-memory relationship, as neither the sigma × arousal interaction nor the sigma × arousal × baseline interaction was significant, all *ps* ≥ .066. By contrast, for valence ratings, the sigma × valence rating × baseline interaction was significant, $\chi^2(1) = 11.85$, $p < .001$. Follow-up trend analyses revealed that when baseline temporal memory was low (-1 SD), greater sigma power predicted poorer subsequent temporal memory for events rated as relatively less aversive, $z = -2.76$, $p = .006$, but not for events with mean  or more aversive valence ratings (all *ps* ≥ .270, Fig. S3B).

### Supplementary Tables

| Effect | $\chi^2$ | Slope | SE | 95% CI | *z* | *p* |
|---|---|---|---|---|---|---|
| **Immediate test** | | | | | | |
| Theta power | 0.08 | — | — | — | — | .782 |
| Theta power × Valence | 4.29 | — | — | — | — | **.038** |
| Control-cue theta power | 1.31 | — | — | — | — | .252 |
| Theta slope: Neutral | — | −0.026 | 0.011 | [−0.047, −0.005] | −2.437 | **.015** |
| Theta slope: Aversive | — | −0.007 | 0.013 | [−0.033, 0.019] | −0.530 | .596 |
| **Delayed test** | | | | | | |
| Theta power | 0.15 | — | — | — | — | .694 |
| Theta power × Valence | 0.75 | — | — | — | — | .385 |
| Control-cue theta power | 0.93 | — | — | — | — | .334 |
| Theta slope: Neutral | — | −0.014 | 0.009 | [−0.032, 0.004] | −1.479 | .139 |
| Theta slope: Aversive | — | 0.003 | 0.012 | [−0.020, 0.025] | 0.242 | .808 |

**Table S3.** Associations between cue-elicited theta power and temporal-order memory at immediate and delayed tests, controlling for responses to control cue.

| Effect | $\chi^2$ | Slope | SE | 95% CI | *z* | *p* |
|---|---|---|---|---|---|---|
| **Immediate test** | | | | | | |
| Sigma power | 1.16 | — | — | — | — | .281 |
| Sigma power × Valence | 2.76 | — | — | — | — | **.097** |
| Control-cue sigma power | 1.03 | — | — | — | — | .311 |
| Sigma slope: Neutral | — | −0.022 | 0.009 | [−0.040, −0.003] | −2.318 | **.020** |
| Sigma slope: Aversive | — | 0.005 | 0.014 | [−0.022, 0.031] | 0.362 | .718 |
| **Delayed test** | | | | | | |
| Theta power | 0.20 | — | — | — | — | .694 |
| Sigma power × Valence | 2.08 | — | — | — | — | .385 |
| Control-cue sigma power | 2.61 | — | — | — | — | .106 |
| Sigma slope: Neutral | — | −0.016 | 0.008 | [−0.032, −0.000] | −1.969 | **.049** |
| Sigma slope: Aversive | — | 0.011 | 0.012 | [−0.012, 0.034] | 0.950 | .342 |

**Table S4.** Associations between cue-elicited sigma power and temporal-order memory at immediate and delayed tests, controlling for responses to control cues.

| Characteristics | Mean | SD |
|---|---|---|
| TIB | 481.73 | 8.31 |
| SPT | 464.97 | 21.10 |
| WASO | 59.66 | 31.58 |
| TST | 405.31 | 35.04 |
| N1 | 20.31 | 8.11 |
| N2 | 205.45 | 34.26 |
| N3 | 92.26 | 27.10 |
| REM | 87.29 | 18.94 |
| NREM | 318.02 | 32.67 |
| SOL | 15.42 | 17.60 |
| Lat_N1 | 28.27 | 44.44 |
| Lat_N2 | 21.35 | 19.07 |
| Lat_N3 | 31.81 | 22.31 |
| Lat_REM | 114.16 | 47.35 |
| %N1 | 5.01 | 1.98 |
| %N2 | 50.81 | 8.03 |
| %N3 | 22.63 | 5.82 |
| %REM | 21.54 | 4.48 |
| %NREM | 78.46 | 4.48 |
| SE | 84.13 | 7.01 |
| SME | 87.19 | 6.65 |

**Table S5.** Descriptive statistics of sleep architecture. Values represent means and standard deviations (SDs) across all participants (N = 31). All duration measures are reported in minutes; percentage measures reflect the proportion of total sleep time (TST) spent in each sleep stage. TIB, time in bed; SPT, sleep period time; WASO, wake after sleep onset; TST, total sleep time; N1/N2/N3, non-REM sleep stages 1–3; REM, rapid eye movement sleep; NREM, non-REM sleep (N1 + N2 + N3); SOL, sleep onset latency; Lat_N1/N2/N3/REM, latency to the first epoch of each stage from sleep onset; %N1/N2/N3/REM/NREM, percentage of TST in each stage; SE, sleep efficiency (TST/TIB × 100); SME, sleep maintenance efficiency (TST/SPT × 100).

### Supplementary Figures

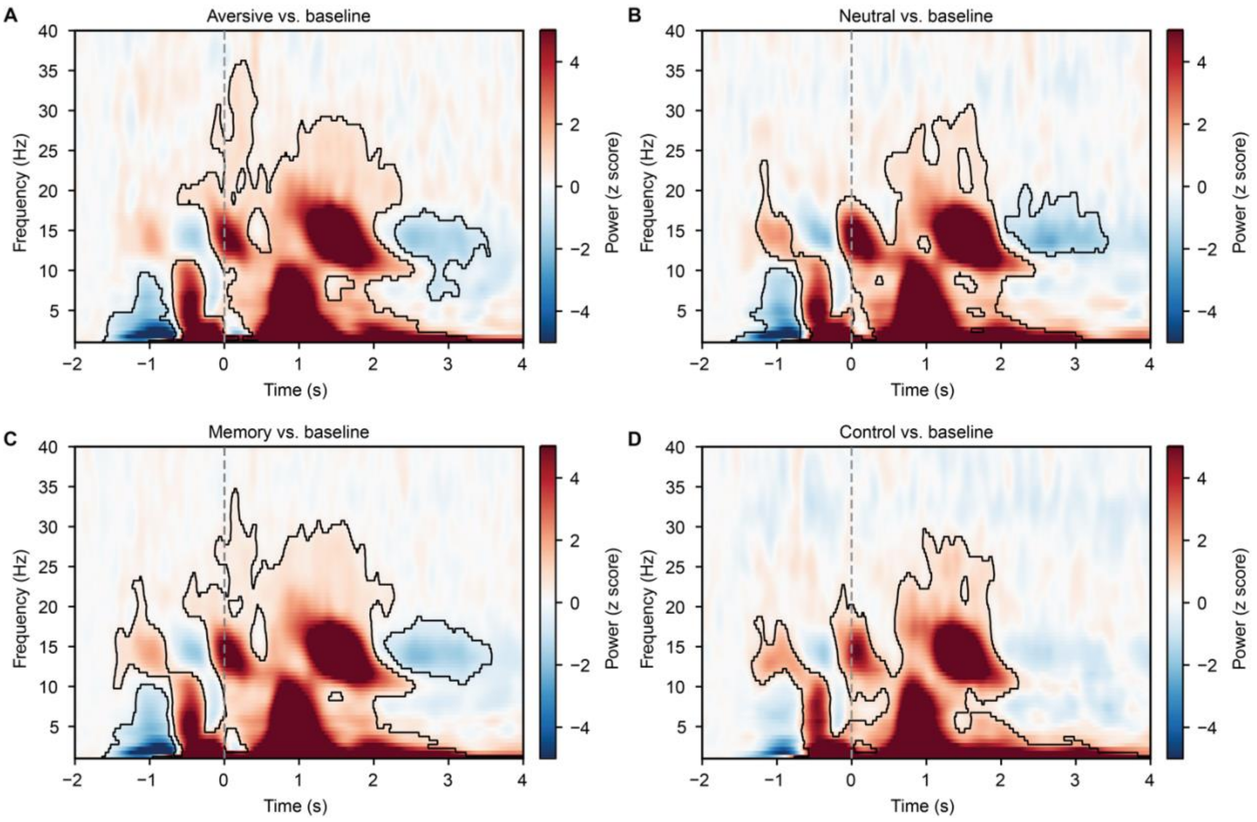


**Fig. S1.** Cue-elicited EEG power changes relative to a pre-cue baseline. Time-frequency representations are shown separately for aversive cues, neutral cues, memory cues (collapsed across aversive and neutral conditions), and control cues. Power values were baseline-corrected using the −2.0 to −1.5 s pre-cue interval. As cues were delivered via a closed-loop procedure time-locked to the detected peak of the slow oscillation, this pre-cue window was chosen to capture spontaneous EEG activity surrounding cue onset ($p_{\text{cluster}} < .05$).

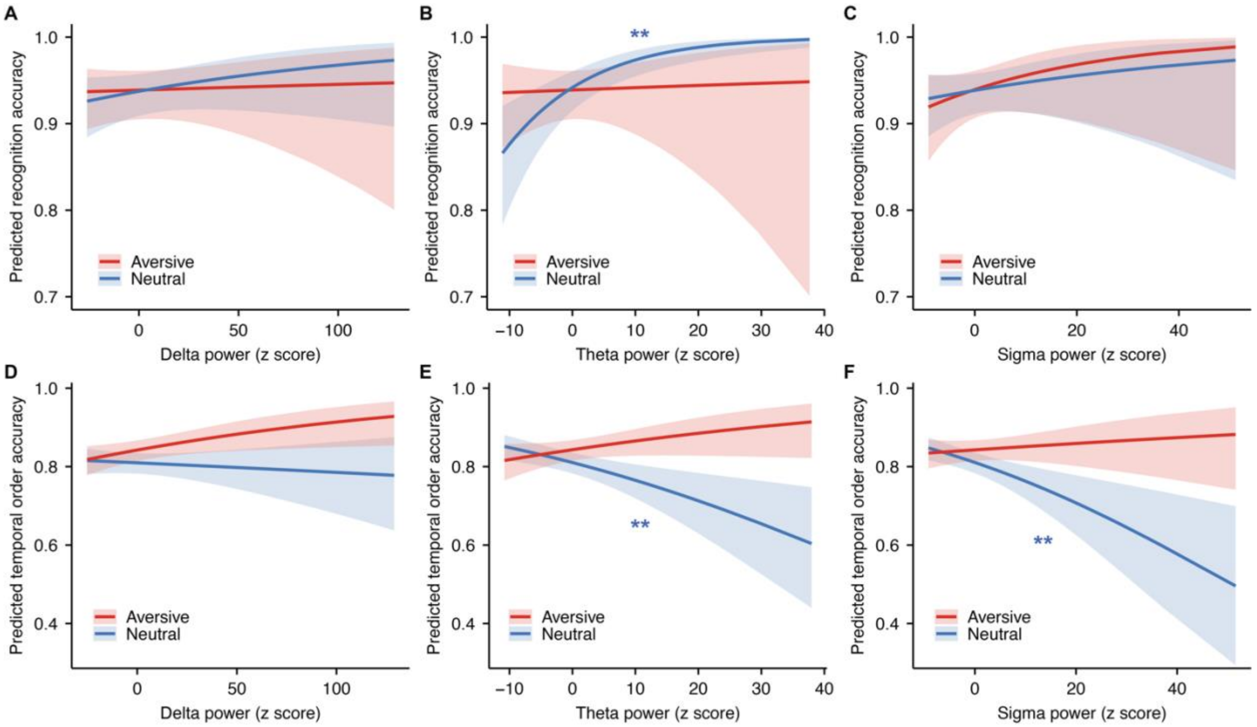


**Fig. S2.** Band-specific EEG predictors of content and temporal-order memory. For content memory, theta power interacted with event valence and boundary position, $\chi^2(1) = 3.93$, $p = .048$. Greater theta power predicted higher recognition accuracy specifically for neutral post-boundary items ($z = 2.65$, $p = .008$), but not for neutral pre-boundary or aversive items (all $ps \geq .422$). Delta and sigma power were not reliably associated with recognition accuracy (all $ps \geq .010$). For temporal order memory, both theta and sigma power interacted with event valence (theta: $\chi^2(1) = 4.46$, $p = .035$; sigma: $\chi^2(1) = 4.83$, $p = .028$). Greater theta and sigma power predicted poorer temporal order accuracy for neutral events (theta: $z = -2.83$, $p = .005$; sigma: $z = -2.91$, $p = .004$), but not for aversive events (both $ps \geq .361$). Delta power showed no reliable valence-dependent association with temporal-order memory, $\chi^2(1) = 2.54$, $p = .111$.

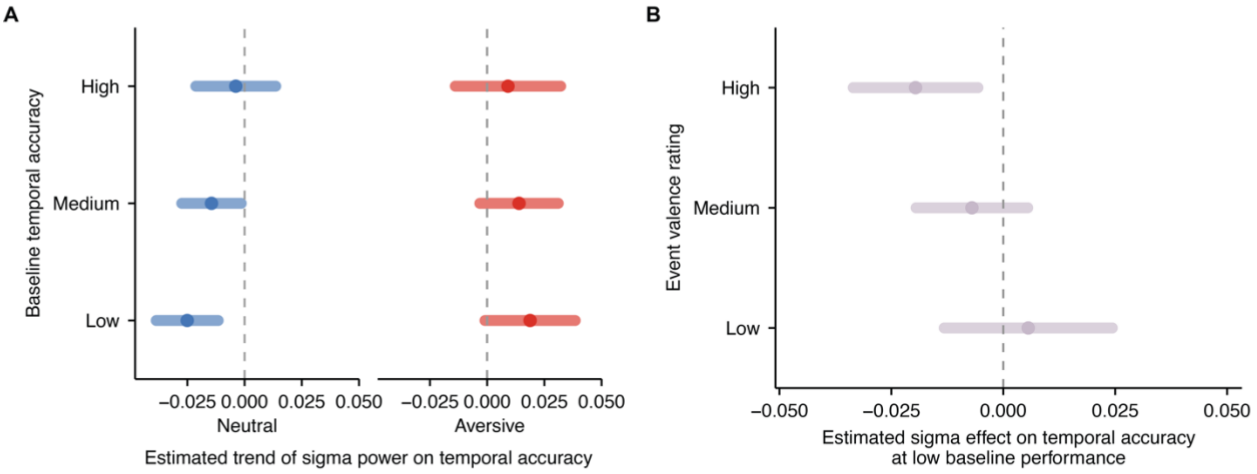


**Fig. S3.** Cue-elicited sigma power negatively predicted temporal order accuracy for neutral events. (A) This sigma–memory relationship was most pronounced when pre-sleep temporal memory performance was low. (B) The sigma–memory relationship was also more pronounced for events rated as less aversive.

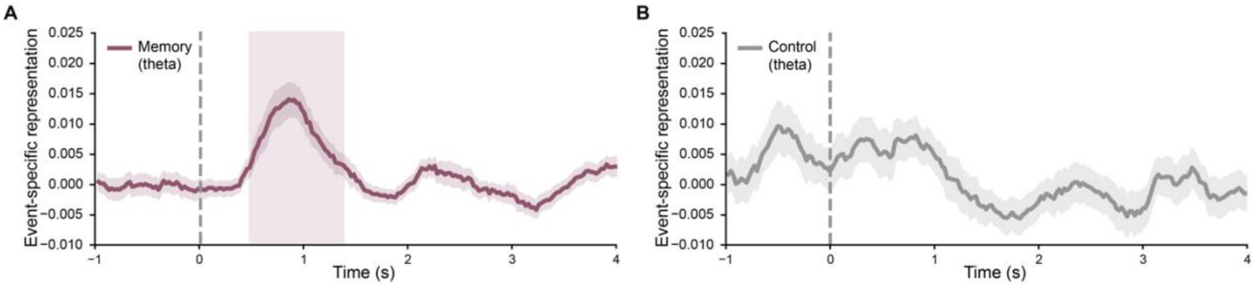


**Fig. S4.** Event-specific representations following strict trial-pair balancing. Trial-pair counts were jointly matched across the memory and control conditions, while maintaining equal numbers of within-event (WE) and between-event (BE) pairs within each condition. Theta-band event-specific representations (WE – BE) remained robust for memory cues (473-1,372 ms, $p_{cluster} < .001$), but were absent for control cues, closely reproducing the significant time window observed in the primary analysis. These results indicate that the observed effect was not attributable to unequal trial-pair counts.
